# HLA alleles and haplotype distribution across Russian population groups

**DOI:** 10.64898/2026.02.25.707989

**Authors:** Varvara Kucherenko, Natalia Doroschuk, Elizaveta Sarygina, Olesya Sagaydak, Viktor Bogdanov, Olga Mityaeva, Julia Krupinova, Mary Woroncow, Eugene Albert, Pavel Volchkov

## Abstract

HLA loci are highly polymorphic genome regions, with allele frequencies varying significantly across different populations. Population HLA frequency databases may contain biases and make cross-study comparison complicated due to varying data curation protocols, genotyping methodologies, resolution, and inconsistencies in the selection criteria for population samples.

This study presents HLA allele frequencies of class I (HLA-A, -B, -C) and class II (HLA-DRB1, -DQB1, -DQA1) as well as their combined haplotypes obtained from over 18,000 whole genome sequencing samples of the Russian population. Cohort was stratified based on PCA and admixture components providing frequencies for 14 different ethnic groups. For 12 groups cohort size allowed us to reach average saturation of 96% of allele frequencies in groups. Moreover, we demonstrated the utility of composed statistics for disease populational study using type 1 diabetes (T1D) as an example. Populations with similar aggregated genetic risk for T1D demonstrated substantial differences in frequencies of risk and protective HLA alleles. Obtained frequency data was made publicly available through the Allele Frequency Net Database improving previously sparse coverage in HLA frequencies data for east Europe and north Asia regions.

## Introduction

Human leukocyte antigen (HLA) diversity, while essential for immunity, creates a major challenge for donor-recipient matching in transplantation. Currently the most efficient way to tackle haplotype matching problem is a construction of a large, usually nation-wide register of HLA haplotypes. Efficient construction of such registries require a good understanding of population-specific variations in HLA haplotype composition to ensure the appropriate coverage for diverse ethnic groups(1). Core clinical typing focuses on HLA-A, -B, -C, -DRB1, -DQB1, - DPB1 and mostly defines transplantation outcome. Extension to -DRB3/4/5 and -DPA1 HLA typing helps reduce the risk of graft failure and allows effective immunosuppression management (2–4).

Population specific HLA haplotype information is aggregated in databases such as Allele Frequency Net Database AFND (5), iHLA-net (6), PGG.MHC (7). AFND provides the most extensive datasets, yet its utility is frequently limited by inconsistencies and variable data quality. In contrast, iHLA-net offers high standards for HLA allele frequencies validation (8,9). For example, it requires the submission of a standardized data format, a detailed description of the allele and haplotype frequency calculation methods, and the inclusion of HWE tests (8). The database is constrained by a small sample size and limited region representation. Meanwhile, PGG.MHC predominantly focuses on East Asian populations: Han Chinese population entry currently has 44523 sample size. Although the database has a methodological description of data processing, it has no explicit data quality stratification (7).

Russia’s territory encompasses a wide range of ethniсities, including ethnic groups from the Caucasus region, Finno-Ugric, indigenous Siberian, Slavic, and Turkic populations. This significant diversity makes the study of HLA variation in the region particularly valuable for both medical and anthropological research. However, comprehensive data on HLA allele and haplotype frequencies across Russia’s distinct ethnic groups remain limited primarily because of heterogeneity of data, small sample sizes, and low resolution. For instance, the iHLA-net records from the Russian region comprise only 3 entries derived from the International HLA & Immunogenetics Workshop, and access to these data is restricted and requires prior permission (10),(11). PGG.MHC includes several (16) Russian populations represented, primarily derived from the Simons Genome Diversity Project, with a median sample size of two individuals (7),(12). AFND has the most extensive population set from the Russian region (53) including Bashkirs, Butyats, Khanty, Mansi, Tatars, Tuvans, etc. Still, some of the ethnicities are underrepresented or entirely absent. For example, the median sample size for Russian AFND entries is only 80. Furthermore, Yakut HLA frequencies are not yet represented in the database.

This study investigates the distribution of HLA class I (HLA-A, -B, -C) and class II (HLA-DRB1, -DQB1, -DQA1) alleles and haplotypes across diverse Russian populations, focusing on key loci with clinical significance. Using whole-genome sequencing (WGS) data from Bashkirs, Chechens, Russians, Tatars, Yakuts, and other groups, we aim to expand current HLA databases by incorporating a large, uniformly processed cohort of high-quality samples. The work explores HLA diversity within and between populations and identifies novel alleles that can refine our understanding of genetic variation across Russia. Ultimately, this research seeks to advance knowledge of the country’s immunogenetic landscape and to support developments in precision medicine, transplantation registries, and population genetics.

## Methods

### 1. Cohort description

Peripheral venous blood samples were collected from 18,548 participants (44.6% male, 55.4% female) by LCC “Evogen” during routine sequencing conducted between 4 April 2025 and 1 December 2025. All participants were healthy individuals from the Russian population and provided written informed consent.

### 2. Sample preparation and sequencing

DNA extraction was performed by spin column using the Qiagen QIAamp DNA Blood Kit (Cat. No. 51106) from whole blood according to the manufacturer’s protocol. DNA amount was measured fluorometrically with Qubit4 (Thermo Fisher Scientific)/Denovix (DeNovix Inc.). For the subsequent library preparation only genomic DNA of high quality (OD260/OD280 = 1.8–2.0, OD260/OD230 > 2.0) was used. Library preparation was performed with a PCR-free enzyme fragmentation protocol (MGIEasy FS PCR-Free DNA Library Prep Set, Cat. No. 1000013455) using 800–1,200 ng gDNA. The distribution of insert size was 400–600 bp. WGS library preparation was performed both manually and automatically.

Whole genome sequencing was performed using DNBSEQ-G400 (MGI Tech Co., Ltd.) with FCL PE150 (cat. no. 1000012555), FCL PE200 (cat. no. 1000013858), and DNBSEQ-T7, according to the manufacturer’s protocol.

### 3. WGS processing

Raw FASTQ files were processed using a modified version of the Broad Institute’s Whole Genome Germline Single Sample pipeline (13) for quality control, mapping and variant calling. A key modification involved replacing BWA-MEM2 with Minimap2 (14) to optimize alignment step. Subsequent analyses included population structure estimates with principal component analysis (PCA) and ADMIXTURE v.1.3.0 (15) under unsupervised mode, NGS HLA typing conducted with HLA-HD (IMGT release 3.53.0) (16) and T1K (17), HLA Haplotype inference performed with Hapl-o-Mat v.1.2.2 (18).

### 4. Ethnic annotation and clustering

We performed genetics-based population labeling using a reference panel of 1,718 samples from previously published Russian population genetic studies (Table S1) (19–23). 38,316 SNP shared across different technologies in these datasets were retained. Reference samples were clustered in two steps: first, DBSCAN (scikit-learn DBSCAN; eps=0.11) on the first two principal components identified 40 clusters; second, agglomerative clustering (scikit-learn AgglomerativeClustering; distance_threshold = 0.1) of cluster centroids refined these to 19 final clusters (Fig. S1). Thresholds were optimized using silhouette scoring.

The resulting reference clusters (paper-based) were used to fit a Random Forest classifier (RandomForestClassifier module, scikit-learn) and assign 18548 individuals from the target dataset to these paper-based groupings. Each cluster was annotated based on its predominant population representatives.

### 5. ADMIXTURE decomposition

Genetic ancestry and population structure were further characterized using ADMIXTURE v1.3.0 (15), employing the same variant set utilized for PCA. The optimal value for K (ranging from K=2 to K=10) was determined by minimizing the five-fold cross-validation error.

### 6. HLA haplotype inference

Haplotypes were inferred using Hapl-o-mat (18). The analyzed haplotype blocks, primarily HLA A B DRB1, and extended versions such as HLA□A□B□C□DRB1 or HLA□A□B□C□DRB1□DQ- are defined by clinical guidelines and published evidence (24–27).

### 7. External validation cohorts and correlation analysis

The validation cohorts were obtained from the AFND (5) and are described in Table S2.

To maximize the utilization of heterogeneous data from the AFND, we performed correlation analyses across all available resolutions and typing methods. This approach was chosen to prevent the significant data loss as a result of restricting the analysis to a single resolution or protocol. We also restricted our analysis to datasets with a sample size greater than or equal to 100. For the comparison of haplotype frequencies we selected AFND entries that exhibited the highest allele frequency concordance with the current study clusters and for which haplotype data were available (N haplotypes > 5).

Pairwise correlation coefficients were calculated between the study-derived population cluster and each corresponding reference population available in the AFND. P-values were adjusted using the Bonferroni correction.

### 8. Analysis of allele richness and allele/haplotype frequency distributions

The proportion of distinct alleles detected was estimated as the ratio of observed allele number to the Chao1 estimator (28) per locus, which infers unseen alleles from the frequency of singletons and doubletons.

Cumulative frequency plots were constructed for each locus to estimate genetic diversity. Alleles/haplotypes were ranked by descending frequency and summed. The resulting sigmoidal curve shows the distribution: a steep rise indicates dominance by a few common alleles, while a gradual slope reflects more even frequencies. By overlaying these curves, we visually assessed similarity between populations.

To balance sample sizes, each cluster was subsampled 10 times to match the smallest group. For cumulative haplotype frequencies, however, the entire sample set was used, as sample size critically impacts Hapl-o-Mat estimates and haplotypes are generally more diverse compared to alleles.

### 9. Polygenic risk scores (PRS) calculation and ethnic HLA associations

Individual PRS were computed as the sum of allele dosages weighted by effect sizes (β) from PGS Catalog (29) summary statistics. Pairwise Mann-Whitney U tests were used to compare the distribution of the Type 1 Diabetes PRS across study populations. P-values were adjusted using the Bonferroni correction.

Using PRS summary weights, T1D haplotypes were designated as risk (positive weight) or protective (negative weight). Their population-specific frequencies were min max scaled per haplotype, then visualized and hierarchically clustered via seaborn.clustermap with default parameters.

## Results

### 1. Ethnic stratification of the cohort revealed 14 distinct clusters showing consistent admixture patterns

The study was done on an ethnically diverse cohort of 18,548 whole-genome sequencing (WGS) samples at 30x coverage. To characterize HLA distribution across different populations we started with characterisation of the cohort population structure. Genetic clusterisation (see Methods) produced 14 distinct ethnic clusters.(Table 1, Fig. 1A).

**Figure. 1.**
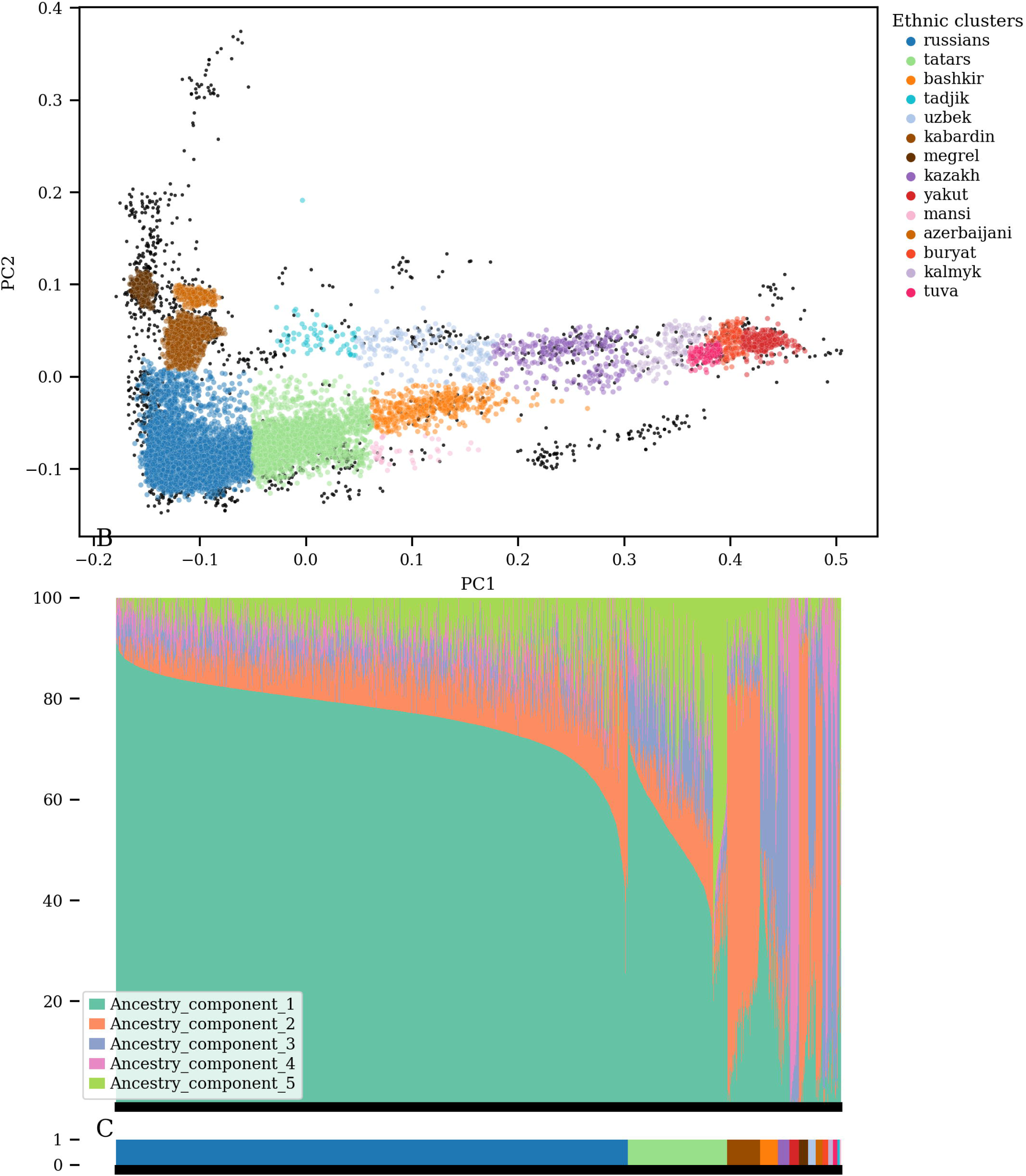
Population structure of the study cohort. A: PCA projection of the cohort; colors indicate population clusters, with reference ethnic samples shown in gray. B ADMIXTURE analysis of ancestry components, where each bar represents an individual sample decomposed into ancestral components contributions. C. Colors in the bar chart correspond to the population cluster legend in the PCA plot.

**Table. 1.** Population cluster sizes.

| Cluster | Size |
| --- | --- |
| russians | 13147 |
| tatars | 2512 |
| kabardin | 834 |
| bashkir | 442 |
| kazakh | 299 |
| yakut | 235 |
| megrel | 230 |
| uzbek | 198 |
| azerbaijani | 170 |
| buryat | 149 |
| kalmyk | 120 |
| tuva | 112 |
| tadjik | 62 |
| mansi | 38 |
| **Total** | **18548** |

To better characterise obtained clusters and to connect them with global populations we performed the additional ADMIXTURE analysis. Analysis confirmed internal homogeneity of identified groups with similar components distributions inside individual clusters (Fig. 1B). Cross-population ancestry decomposition revealed varying contributions of major continental ancestries in line with previously published data. Russians, the largest subgroup, was mostly composed of an ancestry component 1, reflecting apparently European ancestral component (mean proportion: 76.6%, SD: 7.8%), with an admixture of ancestry component 2 (mean: 8.5%, SD: 6.7%) that was predominant in Caucasus ethnic clusters (megrel, kabardin, azerbaijani). In contrast, megrel, kabardin, azerbaijani groups exhibited a reverse admixture pattern, characterized by a high proportion of ancestry component 2 as dominant component (67.0%-87.9%) combined with significant ancestry component 1 (European) admixture (6.7%-16.2%). This component combination was consistent with expected admixture patterns in European-Caucasus populations (30). “Eastern” clusters: yakut, buryat, tuva were defined by ancestry components 3 and 4. Tatars group was mainly a mixture of ancestry components 1 (European) and 5.

These results are concordant with previous studies: five ancestry components were optimal, and the Russian cluster demonstrated a gradient from Northern European to Eastern patterns (31).

### 2. High correlation of allele frequencies with public data supports ethnic clusterisation and HLA typing approach

To ensure the validity of our HLA-typing results and population stratification, we compared the observed HLA allele frequencies (AF) with those reported in publicly available sources (see Methods). Only 8 out of 14 clusters had a complement dataset in the AFND with a sample size of at least 100 samples.

As expected, the largest clusters, russians and tatars, showed the highest observed pairwise correlation coefficients (Fig.2A, Fig.S2-S13). However, several validation AFND datasets exhibited systemic discordance in rare allele frequencies (AFND Russia Bashkortostan, Tatars; Russia Nizhny Novgorod, Russians; Russia Bashkortostan, Bashkirs; Fig S2, S5, S11). Notably, each of these datasets was generated using NGS platforms, in contrast to the alternative HLA typing methods employed for other populations (Table S2). Despite observed heterogeneity in allele frequency correlations across study populations (SD: 0.04–0.29) all clusters maintained a mean Pearson correlation coefficient exceeding 0.8 (Table S3).

**Figure 2.**
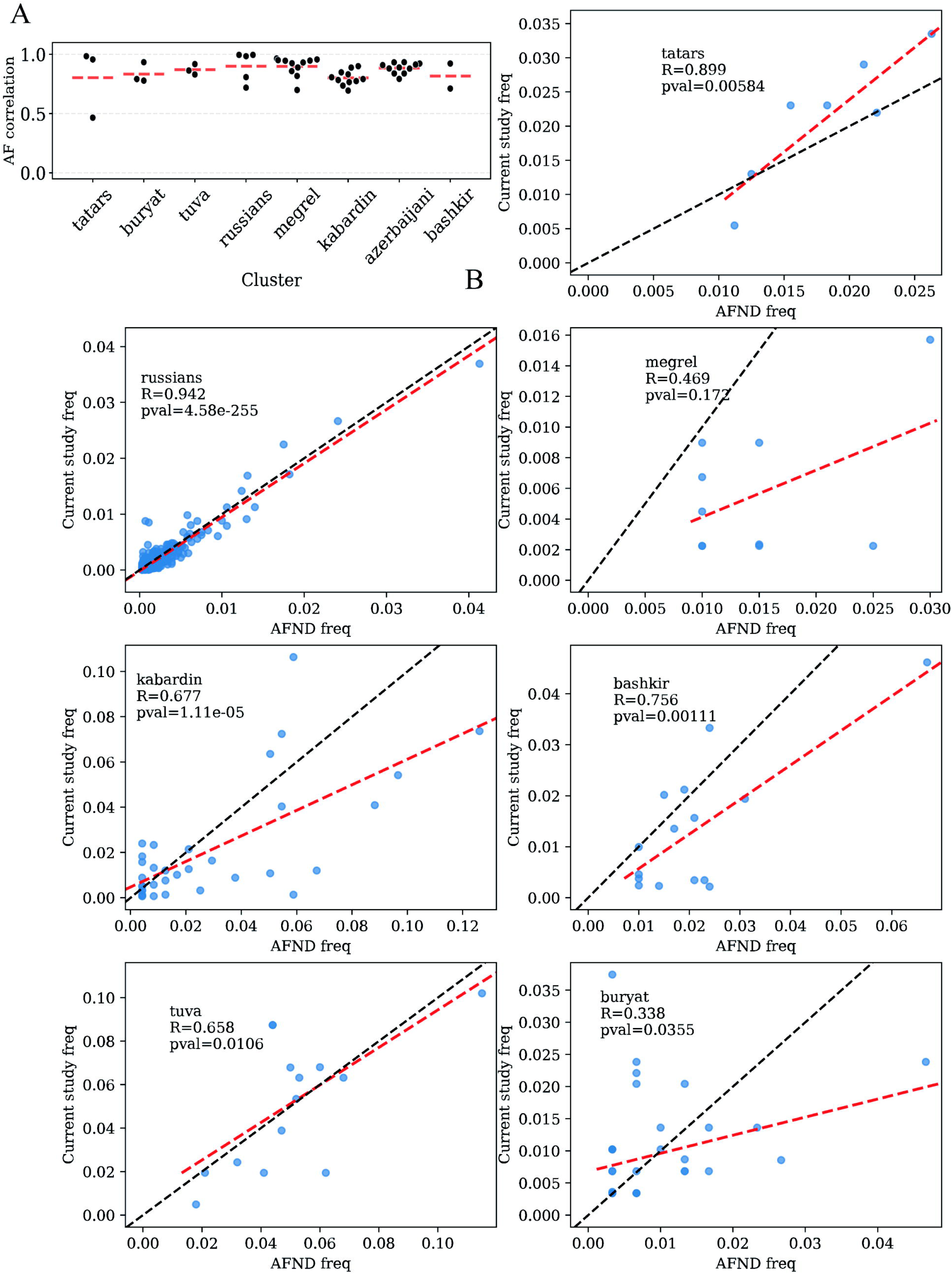
Concordance between the current study and AFND Statistics. A. Allele Frequency correlation of the current study clusters and multiple AFND populations. Each dot represents a pairwise correlation of the current study population cluster AF vs AFND AF dataset. Mean AF correlation per population cluster is shown by red dashed lines; B. Frequency correlation of the current study cluster haplotypes and AFND population haplotypes, with the black dashed line representing the line of perfect haplotype frequency agreement (y=x) and red dashed line showing linear approximation.

We then assessed the reliability of our HLA genotyping data by reconstructing haplotype frequencies and comparing them to reference data from the AFND. Due to sparse haplotype records in the AFND compared to allele data (Table S2), haplotype frequency distributions are less consistent across clusters, except for the russians group, which exhibits the highest correlation (R=0.942, Fig.2B) likely due to its homogeneous internal population structure and significantly larger sample size both in the current study and in the AFND. Smaller clusters tend to have fewer haplotype frequency records in AFND and also lack well-matched reference populations, thereby reducing the reliability of correlation estimates.

To summarize, for overlapping populations between our cohort and AFND we saw a high allele frequency correlation which supports the reliability of our data. However, haplotype frequencies show limited concordance. This issue particularly affects small clusters.

### 3. Frequency saturation analysis reveal reduced HLA diversity of the yakut population cluster

Because of the extreme variability of HLA alleles large cohorts are required for diversity characterisation. The Chao1 estimates of allelic richness (Table 2) shows that the genetic diversity within the studied cohort is effectively represented across both the large and comparatively smaller identified clusters. We also analyzed the cumulative frequencies of alleles and haplotypes (Fig. 3). Locus-wise curve matching revealed homogeneity in HLA allele cumulative frequency profiles across most clusters especially when it comes to HLA class II loci. However, the yakut population cluster exhibited notable divergence in HLA-A, HLA-B and HLA-C allele frequencies (Fig. 3A). The buryat cluster demonstrates a similar to yakuts pattern, consistent with its Asian origin.

**Figure 3.**
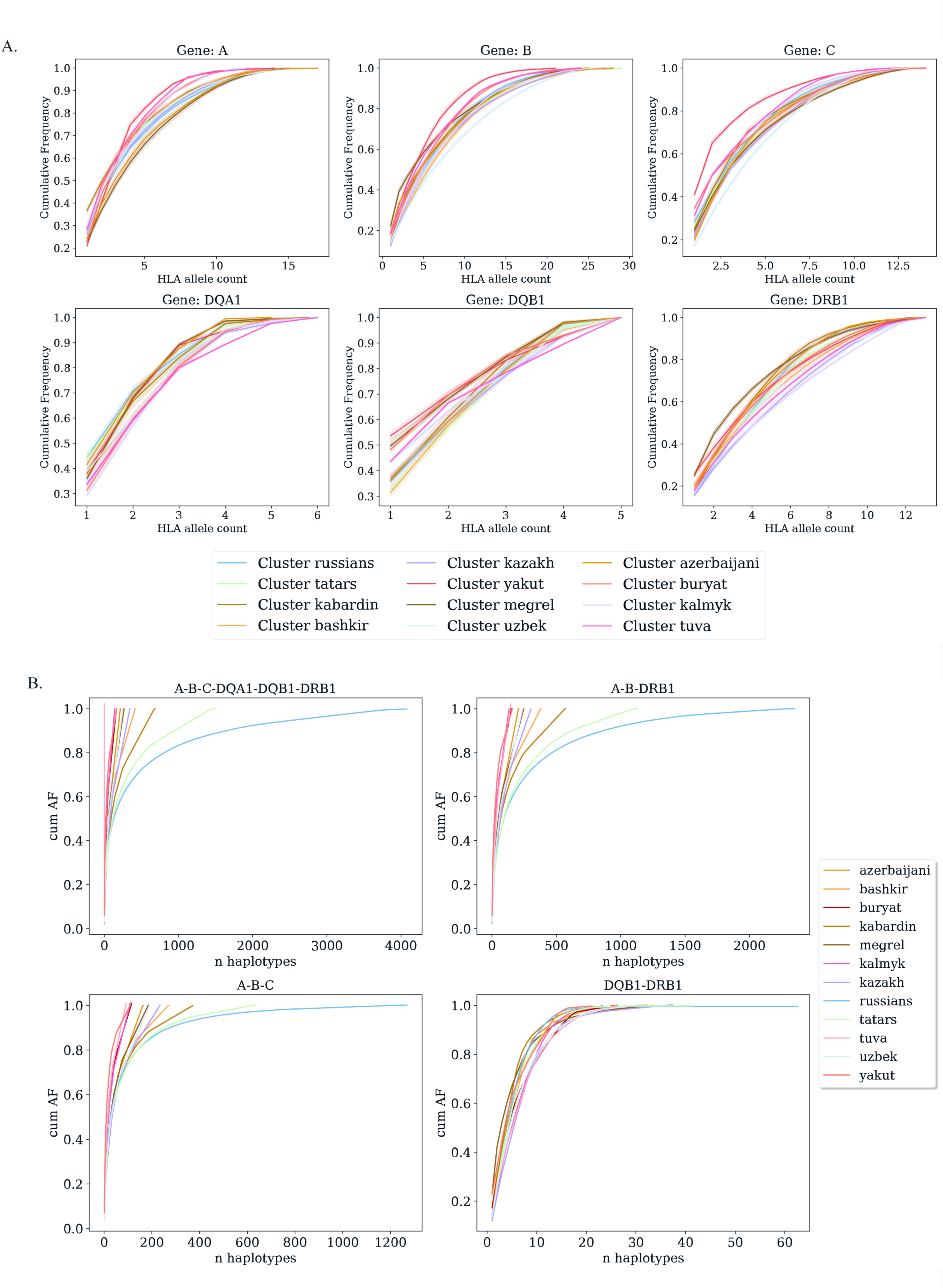
Cumulative HLA allele and haplotype frequencies. A. Cumulative HLA population allele frequencies per HLA-loci. Note prominent red line representing the yakut genetic cluster for genes B and C. B. Cumulative frequency plots for haplotypes of various population clusters. A-B-C and DQB1-DRB1 haplotypes, though not clinically relevant, were considered for HLA class I and HLA class II diversity comparison.

**Table. 2.** The Chao1 estimates of allelic richness in the study cohort.

| Cluster | Mean | SD |
| --- | --- | --- |
| azerbaijani | 0.913 | 0.06 |
| bashkir | 0.965 | 0.033 |
| buryat | 0.934 | 0.066 |
| kabardin | 0.973 | 0.017 |
| kalmyk | 0.943 | 0.057 |
| kazakh | 0.962 | 0.042 |
| megrel | 0.962 | 0.033 |
| russians | 0.994 | 0.004 |
| tatars | 0.987 | 0.008 |
| tuva | 0.976 | 0.019 |
| uzbek | 0.937 | 0.051 |
| yakut | 0.972 | 0.035 |

**Table. 3.** Exonic genetic variants in the mansi population cluster. The REF column refers to a reference allele from the IMGT HLA allele database. POS column shows coordinate in concatenated exon reference sequence.

| Allele | POS | REF | ALT | n reads support alt | n of reads cover this position | n of reads that uniquely aligned support ralt | the coordinate with respect to the input reference |
| --- | --- | --- | --- | --- | --- | --- | --- |
| HLA-B*35:500 | 617 | G | C | 10 | 12 | 3 | 1040 |
| HLA-C*03:524 | 109 | C | T | 7 | 9 | 3 | 288 |

Next we assessed whether clinically relevant haplotypes show cross-population variability. Additionally we included separate HLA class I and II haplotypes in analysis to estimate class contribution to population divergence (Fig.3B). Since the number of IMGT HLA (32) entries for class I is twice greater than for class II, HLA class I exhibits higher diversity than class II. Consequently, a frequency plateau is not detectable for class I-containing haplotypes (A-B-C, A-B-C-DQA1-DQB1-DRB1,A-B-DRB1) in most populations due to insufficient sampling (Fig.3B). This effect is roughly observable in the russians and tatars clusters only, where sample sizes were adequate to capture the asymptotic stabilization of the curve.

Thus, the allele frequency data can be considered representative. That allows us to infer a more modest allele diversity among yakut samples. Data on the frequencies of clinically relevant haplotypes remain insufficient for robust inter-population analysis.

### 4. Type one diabetes-associated HLA haplotypes differentially represented across populations and shape polygenic risk score distribution

HLA alleles represent one of the strongest genetic determinants of immune regulation and disease susceptibility. Thus population HLA frequency differences should be reflected in the mean population risks of immune disorders. The substantial contribution of HLA loci to these risk estimates is particularly evident in type 1 diabetes mellitus (T1D).

Firstly, we calculated PRS for type 1 diabetes (T1D) PGS000024 (28) in our cohort. The score PGS000024 was chosen because it explicitly includes HLA alleles and their interaction instead of proxy SNPs. Identified previously ethnic groups demonstrated 29 statistically significant pairwise differences based on mean PRS. (Fig.4A, Table S4). The most explicit PRS distinction was observed between European and Caucasus ethnic groups, compared to East Asian groups; comprising tuva, yakut, and buryat clusters, which aligns with established patterns of HLA allele frequencies (32).

**Fig. 4.**
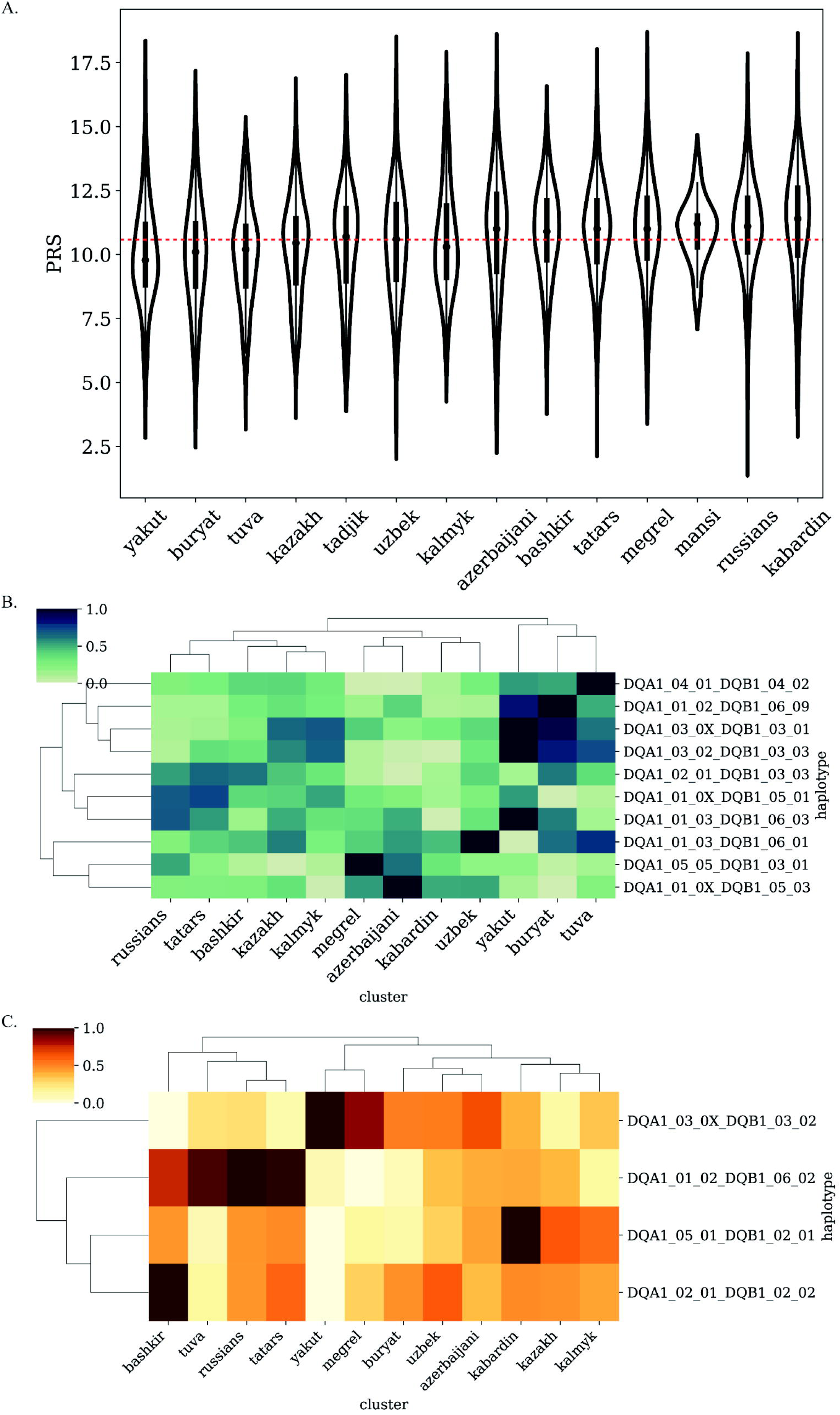
Distribution of T1D genetic risk score and associated HLA haplotypes across populations. A. PRS T1D distributions across population clusters. Dashed lines show mean values of T1D PRS across all populations; B. Frequencies of T1D-protective HLA haplotypes (negative PRS weights), min-max scaled per haplotype across population clusters.; C. Frequencies of T1D-risk HLA haplotypes (positive PRS weights), min-max scaled per haplotype across population clusters.

Secondly, we estimated the contribution of different HLA alleles into the PRS of different ethnic clusters. PGS000024 accounts for both risk and protective HLA alleles. Protective HLA haplotype distribution across ethnic clusters demonstrated the same differences between large european and asian groups as the overall PRS (Fig. 4B). The “asian” clusters (yakut, tuvan, buryat), which exhibit the lowest aggregate risk scores, form a distinct group defined by a high prevalence of established protective haplotypes. In contrast, T1D risk haplotypes demonstrate different distributions within large ethnic groups (Fig.4C). While yakuts and buryats remain together, tuva cluster is closer to russians, bashkirs, and tatars. Kabardin group, characterized by the highest mean PRS, demonstrates a highly expressed risk haplotype DQA1*05:01∼DQB1*02:01, which separates this group from other lower-risk populations. Overall differences in the HLA allele frequencies occasionally result in a similar PRS score which is based on a different set of risk and protective alleles. For example, megrel and tatars, while exhibit statistically indistinguishable PRS, have completely different HLA profiles. Within the protective haplotype set, the megrel cluster is characterized by high-frequency haplotypes DQA1*05:05∼DQB1*03:01 and DQA1*01:0X∼DQB1*05:03. In contrast, the tatar profile is dominated by DQA1*01:0X∼DQB1*05:01 and DQA1*02:01∼DQB1*03:03. In case of risk haplotypes megrel group show a pronounced prevalence of DQA1*03:0X∼DQB1*03:02, that brings this cluster closer to yakut profiles than to tatars (DQA1*01:02∼DQB1*06:02 is predominant).

These results indicate that identical aggregate risk estimates can mask a potentially different immunogenetic background specific to population.

### 5. Discovering new HLA alleles in mansi cluster

Small, genetically isolated ethnic groups remain underrepresented in genomic studies, yet they hold significant potential for uncovering novel allelic variants. Leveraging the advantage of a large cohort, we tried to estimate the prevalence of a new allele in a poorly described population. Mansi is a small (around 12000) indigenous Ugric ethnic group living in Western Siberia, Russia, primarily in the Khanty-Mansi Autonomous Okrug–Yugra (34). We applied T1K and identified 232 potentially novel SNPs that passed default filters in a mansi group in our cohort (38 individuals). While the majority of these were variants in pseudogenes, there were 2 previously unreported genetic variants across the target HLA class I and II loci (A, B, C, DQA1, DQB1, DRB1), carried by 2 individuals (Table.3).

Although further validation is required, these findings highlight the presence of population-specific HLA alleles that may reflect unique selective pressures or founder effects within this cluster and gives an opportunity for further expanding the IMGT HLA database.

## Discussion

The present study has provided an overview of HLA allele and haplotype frequencies across different Russian population groups, contributing to the understanding of genetic diversity, population structure, and future disease susceptibility, transplantation medicine, or anthropological studies.

Given the substantial sample size and uniform HLA typing method, our dataset provides broader coverage of Russia’s major populations, including smaller ethnic groups than previously published research, which focuses on individual populations or does not subdivide large multinational cohorts (35–38). Ethnic groups were identified exclusively by genetic clusterisation, which carries inherent advantages and limitations. This methodology is a scalable framework for participant recruitment, which allows researchers to conduct studies prior to resource-consuming ethnographic expeditions. Genetic clusterisation also provides measurable, objective information compared to the self-reporting ethnicity. However, since sample recruitment did not account for genealogy or geographic origin, the resulting groups exhibit admixture of varying origins, potentially blurring the borders of distinct population clusters. Moreover, cluster size disbalance between subpopulations may affect representativeness. Nevertheless, the derived clusters encompass both European-related groups (e.g., russians, kabardin) and Asian-related populations (e.g., yakut, tuva), reflecting the country’s diverse genetic landscape.

Frequency data presented here align well with allele frequency from AFND. Nevertheless, AFND exhibits several notable limitations including modest sample size for some populations and lack of the other populations. Due to these constraints only 8 ethnic clusters out of 14 identified have corresponding AFND entries to compare. Despite good overall correlation, some NGS-derived datasets from the AFND have demonstrated notable discordances (median R=0.71) in reported allele frequencies, which may be due to insufficient quality control of the submitted datasets. For example, a clear data inconsistency is evident in the AFND Russia Bashkortostan, Tatars dataset. The reported frequency for the allele DQB1*03 is only 0.0026. This is highly questionable, because the frequency of the DQB1*03:01 allele in the same AFND dataset is 0.1875. Since the general allele group DQB1*03 must include all its subgroups (e.g. DQB1*03:01, DQB1*03:02), its frequency cannot be lower than that of any one of them. Several populations (e.g., azerbaijani and kabardin clusters) lack specific representative AFND reference data. The DQA1 data for tuva and buryat clusters is constrained to a single available AFND dataset without independent validation from additional reference up-to-date HLA typing data. Such pronounced inconsistencies and data sparsity clearly shows the need for a larger, uniform data source.

We also analysed haplotype concordance. Haplotypes offer greater informational density than single alleles for enhanced diagnostics and population analysis (39,40). However, technological challenges in haplotype phasing have limited the availability of haplotype data compared to genotype data. Insufficient data reliability stems from key constraints: (1) sparser haplotype representation in AFND than allele data, (2) ambiguous AFND inference methodologies, (3) scarce data for Russian ethnic minorities, and (4) phasing biases in EM-based refinement strategies. Hence, meaningful comparative analysis is only relatable for the largest groups in our datasets - russians and tatars. In other cases comparisons reveal only relative concordance. Despite substantially greater sample recruitment compared to previous AFND entries and the ability to capture HLA allele frequency diversity, this study does not capture the full HLA haplotype repertoire for minor ethnic clusters.

Since our data demonstrate a sufficient level of concordance with previous studies while also benefiting from a uniformly typed large ethnic sample and a high-resolution method for processing genetic data, we tried to assess the diversity of alleles and haplotypes in different ethnic groups and estimate inter-population variability. Analysis showed that the yakut cluster exhibits a lower HLA allele diversity relative to other studied clusters. Apparently, HLA haplotypes and alleles show evidence of founder effect followed by genetic drift which is supported by earlier studies of Y-/MT-haplogroups (19,41) and can be an important feature for optimal donor-recipient matching.

What’s even more important for optimal donor-recipient selection is the identification of previously undescribed HLA alleles. This issue is particularly relevant for minority ethnic groups in Russia. Our study provides an example of searching potentially new alleles within a distinct narrow subpopulation of the current cohort. Mansi group (38 samples) exhibited 0.44% of potentially novel allele rate which hardly can be extrapolated to the general population of Mansi but underlines the underrepresentation of minor ethnic groups in public data.

Population characteristics of HLA allele distribution can bias PRS. Statistically significant differences across population clusters were observed in T1D PRS (PGS000024), largely driven by the HLA component. The tuva, buryat, and yakut clusters showed lower average risk than the russians cluster, aligning with regional epidemiology(42). However, the kabardin cluster’s PRS deviated from expected risk despite a typical regional T1D incidence(42). The observed results could stem from both sample composition (genetic vs. geographic clustering) and the acute need of population-dependent distinct score corrections. Importantly, even statistically similar risk scores can result from different underlying HLA haplotype distributions, suggesting possible population-specific etiopathogenetic mechanisms. Thus, describing ethnic variability in HLA haplotypes and implementing population stratification are essential for appropriate PRS implementation.

Thus we’ve created a novel, uniformly processed genomic dataset of HLA allele and haplotype frequencies encompassing a wide spectrum of Russian ethnicities, which has been submitted to AFND. Current study contributes to public genomic databases that have notable deficiency of representative samples and can serve as a potential reference for further population genetics and epidemiology studies.

## Data availability

Allele and haplotype frequency data supporting the findings of this study have been submitted to the AFND (5) and will be made publicly available upon acceptance. Individual-level genetic data are governed by a data use agreement and are not publicly available due to privacy and confidentiality restrictions.

All supplementary tables and figures associated with this paper are provided in the Supplementary Information files.

## Ethics approval

The study was approved by the local ethical committee of the Independent Multidisciplinary Committee on Ethical Review for Clinical Trials (Moscow, Russia) and was performed in accordance with the approved guidelines and the Declaration of Helsinki. Inform consent was obtained from all participants.

## Acknowledgments

Whole genome sequencing and sample collection were performed in the Evogen LLC by laboratory staff (Antonenko A., Aydarova V., Belov R., Bartsis A., Boldireva B., Bykadorov P., Dibirova H., Domoratskaya E., Frolkov A, Hahanova V., Gubona M., Golovanova M., Leonova V., Markov D., Mikhaylov V., Nikitina O., Panferova A., Pudova L., Revkova M., Rodionova D., Safina S., Shichkova A., Sokolova N., Ulanova P., Vasilenko A., Veselova G., Zolotopup A., deputy lab head Krinitsina A., and lab head Belenikin M.).

## Author Contributions

EA conceptualization, supervision, data processing, editing; VK investigation, data processing, writing, editing; ND, ES, OS sample collection and sequencing.

All authors participated in the discussion and reviewed the manuscript.

## Supporting Information

Supplementary Table.S1 Ethnic annotation of 1718 reference samples from publicly available sources

Supplementary Tables and Figures

