## Supplemental Tables and Figures for "HLA alleles and haplotype distribution across Russian population groups"

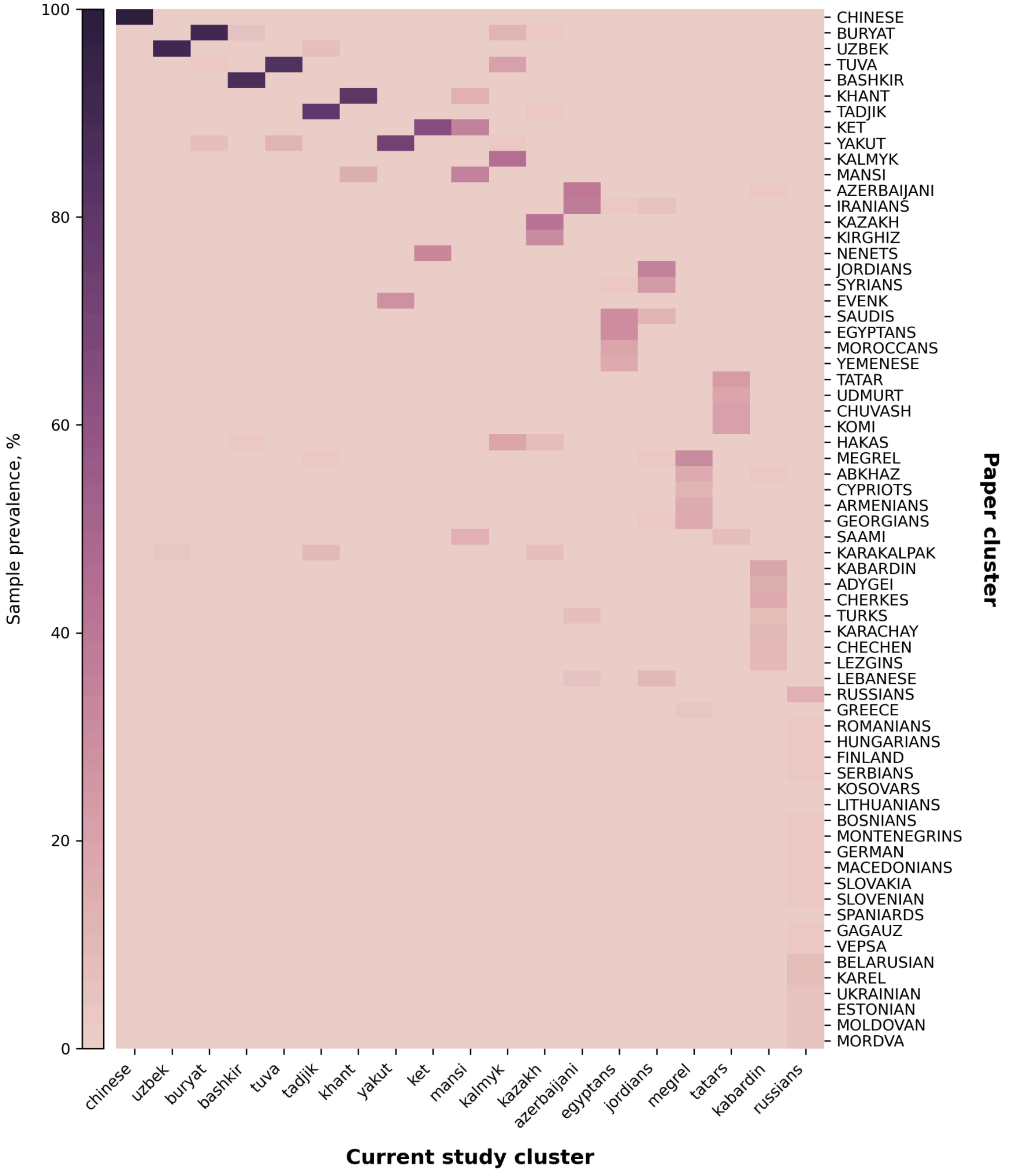


Fig. S1 Distribution of 1718 ethnically-annotated reference samples across current study genetic clusters. Columns represent the genetic clusters identified in the current study. Rows represent the ethnicity labels reported in the source publications. Each cell shows the percentage of samples within a given cluster (column) that were originally annotated with a specific ethnicity (row).

Table.S2. Population clusters of the current study and the corresponding populations from the AFND database. AFND HLA typing methods are NGS (Next Generation Sequencing), SBT (Sequence Based Typing), SSP (Sequence-Specific Primers), SSOP (Sequence-Specific Oligonucleotide Probes). Column AF/Haplotypes denotes if data for HLA alleles or haplotypes frequency is available. The Current Cluster column maps corresponding genetic groupings for direct frequency comparisons. The Resolution column specifies the degree of allelic typing quality in the AFND entry, expressed as the number of digits in the reported HLA genotype resolution.

| AFND population | AFND sample size | AFND HLA typing method | Source | AF/ Haplotypes | Current Study Cluster | Resolution |
|---|---|---|---|---|---|---|
| Russia Bashkortostan, Tatars | 192 | NGS | Bone Marrow Registry | both | tatars | mixed(2-8) |
| Russia Tatars | 355 | SBT | Bone Marrow Registry | both | tatars | 2d |
| Russia South Ural Tatar | 135 | SSP | Anthropology Study | both | tatars | mixed(2-4) |
| Russia Tuva pop 2 | 169 | SSOP | Anthropology Study | both | tuva | mixed(2-6) |
| Russia Transbaikal Territory Buryats | 150 | SSOP | Bone Marrow Registry | both | buryat | mixed(2-4) |
| Russia Northwest pop 2 | 346 | SSP | Controls for Disease Study | AF | russians | 2d |
| Russia Moscow Pop 2 | 2000 | SSP, SSOP | Bone Marrow Registry | both | russians | 2d |
| Russia Moscow | 2650 | SSP, SSOP | Moscow Stem Cell Bank | AF | russians | 2d |
| Russia Nizhny Novgorod, Russians | 1510 | NGS | Bone Marrow Registry | both | russians | mixed(4-8) |
| Armenia combined Regions | 100 | SSP | Bone Marrow Registry | both | megrel | mixed(2-4) |
| Georgia Tbilisi | 109 | SSOP | Anthropology Study | AF | megrel | mixed(2-8) |
| Russia North Ossetian | 127 | SSP, SSOP | Bone Marrow Registry | both | megrel | mixed(2-4) |
| Armenia combined Regions | 100 | SSP | Bone Marrow Registry | both | kabardin | mixed(2-4) |
| Georgia Tbilisi | 109 | SSOP | Anthropology Study | AF | kabardin | mixed(2-8) |
| Russia Bashkortostan, Bashkirs | 120 | NGS | Bone Marrow Registry | both | bashkir | mixed(2-8) |
| Russia South Ural Bashkir | 146 | SSP | Anthropology Study | both | bashkir | mixed(2-4) |
| Russia North Ossetian | 127 | SSP, SSOP | Bone Marrow Registry | both | azerbaijani | mixed(2-4) |

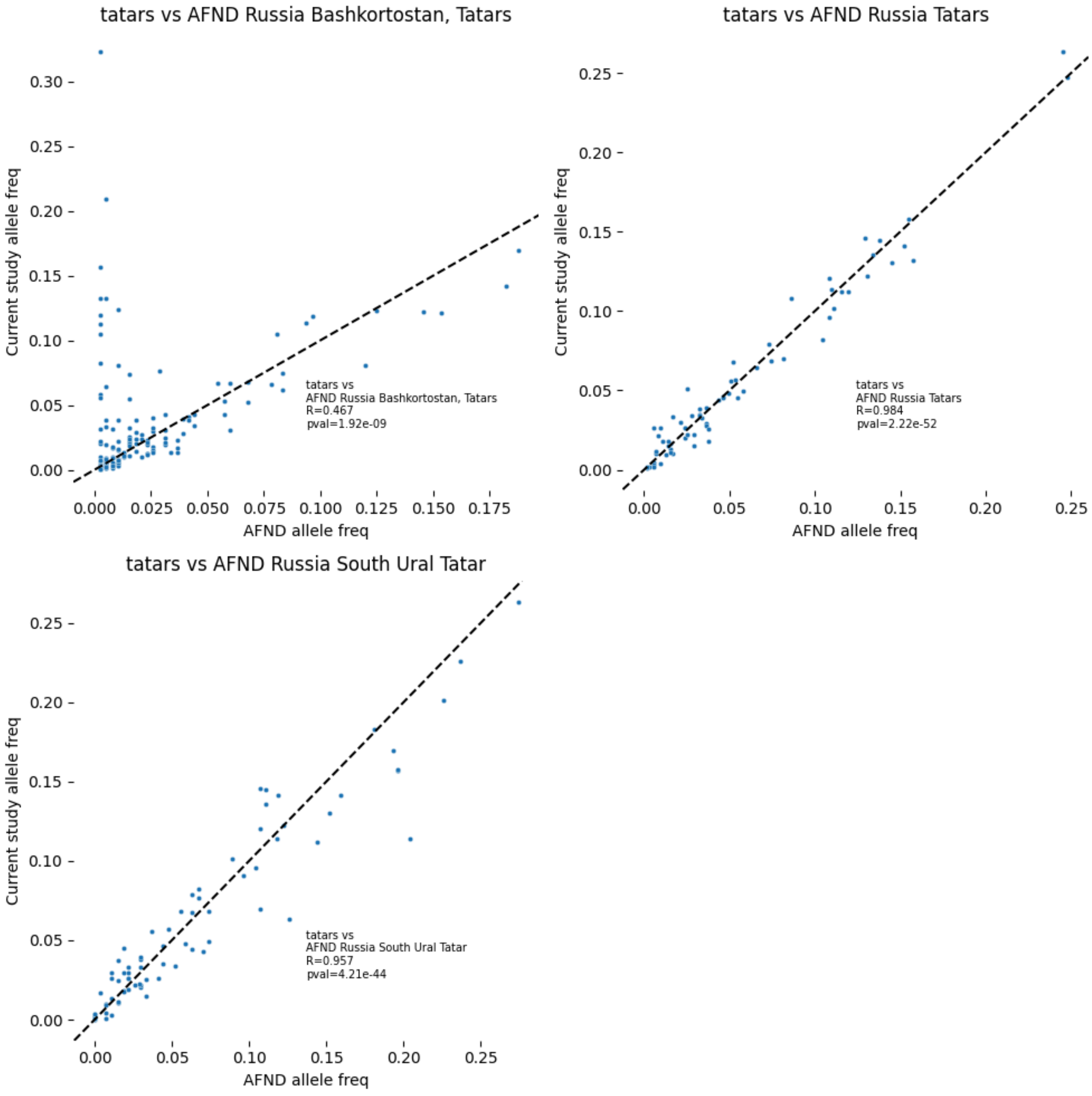


Fig S2. Allele frequency correlation of the current study cluster tatars and the corresponding AFND datasets, with the black dashed line representing the line of perfect haplotype frequency agreement (y=x).

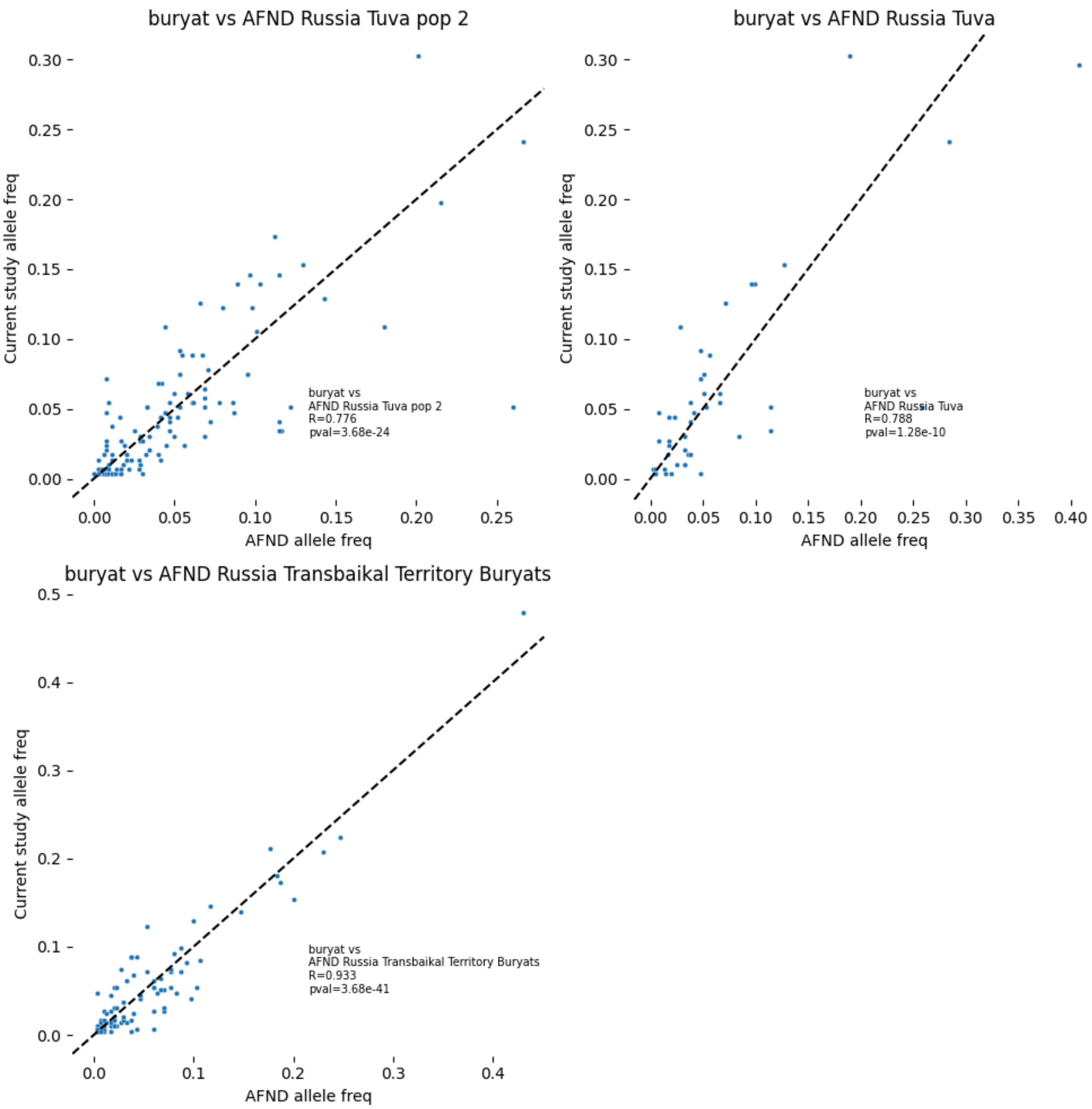


Fig S3. Allele frequency correlation of the current study cluster buryat and the corresponding AFND datasets, with the black dashed line representing the line of perfect haplotype frequency agreement (y=x).

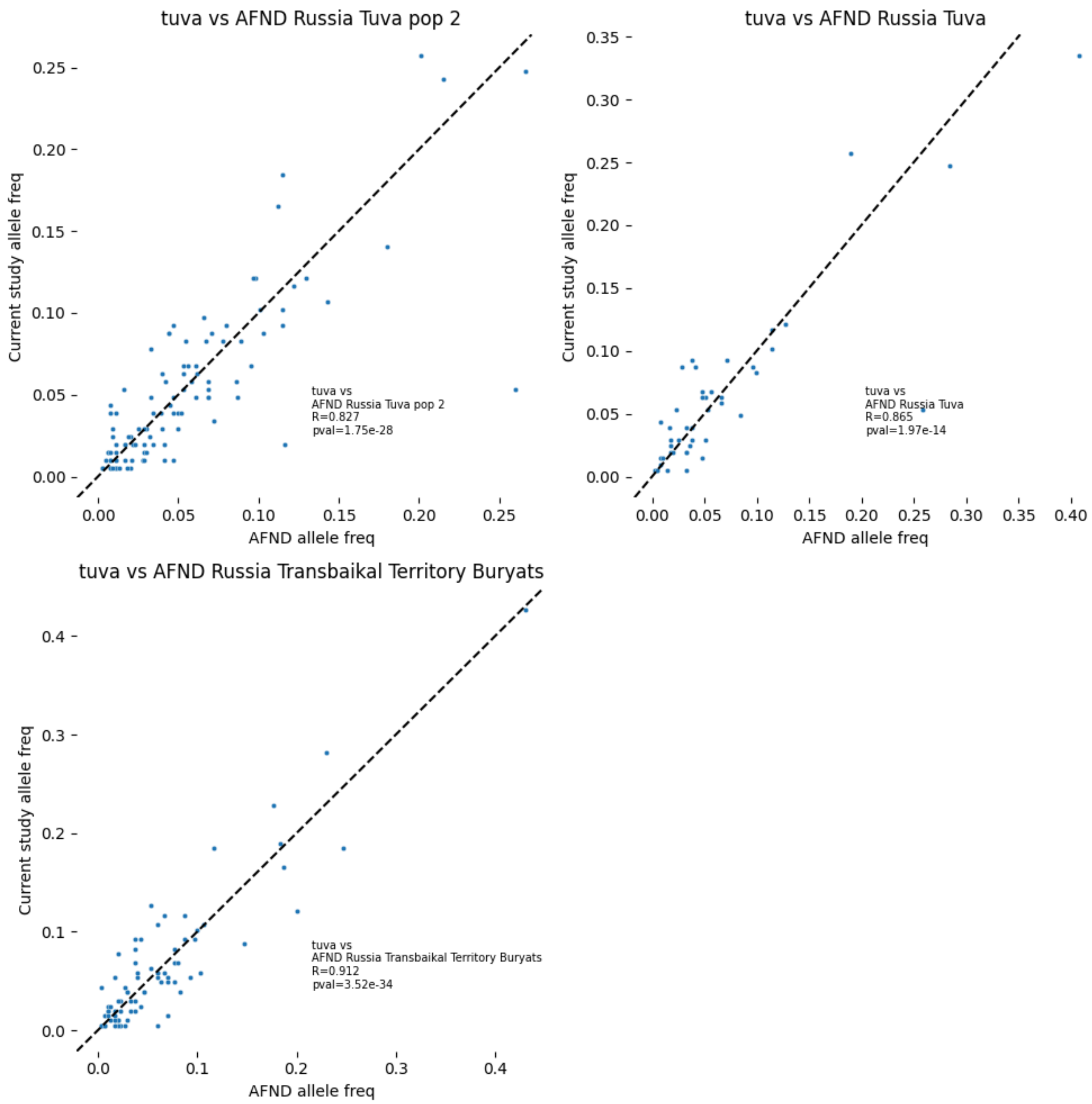


Fig S4. Allele frequency correlation of the current study cluster tuva and the corresponding AFND datasets, with the black dashed line representing the line of perfect haplotype frequency agreement (y=x).

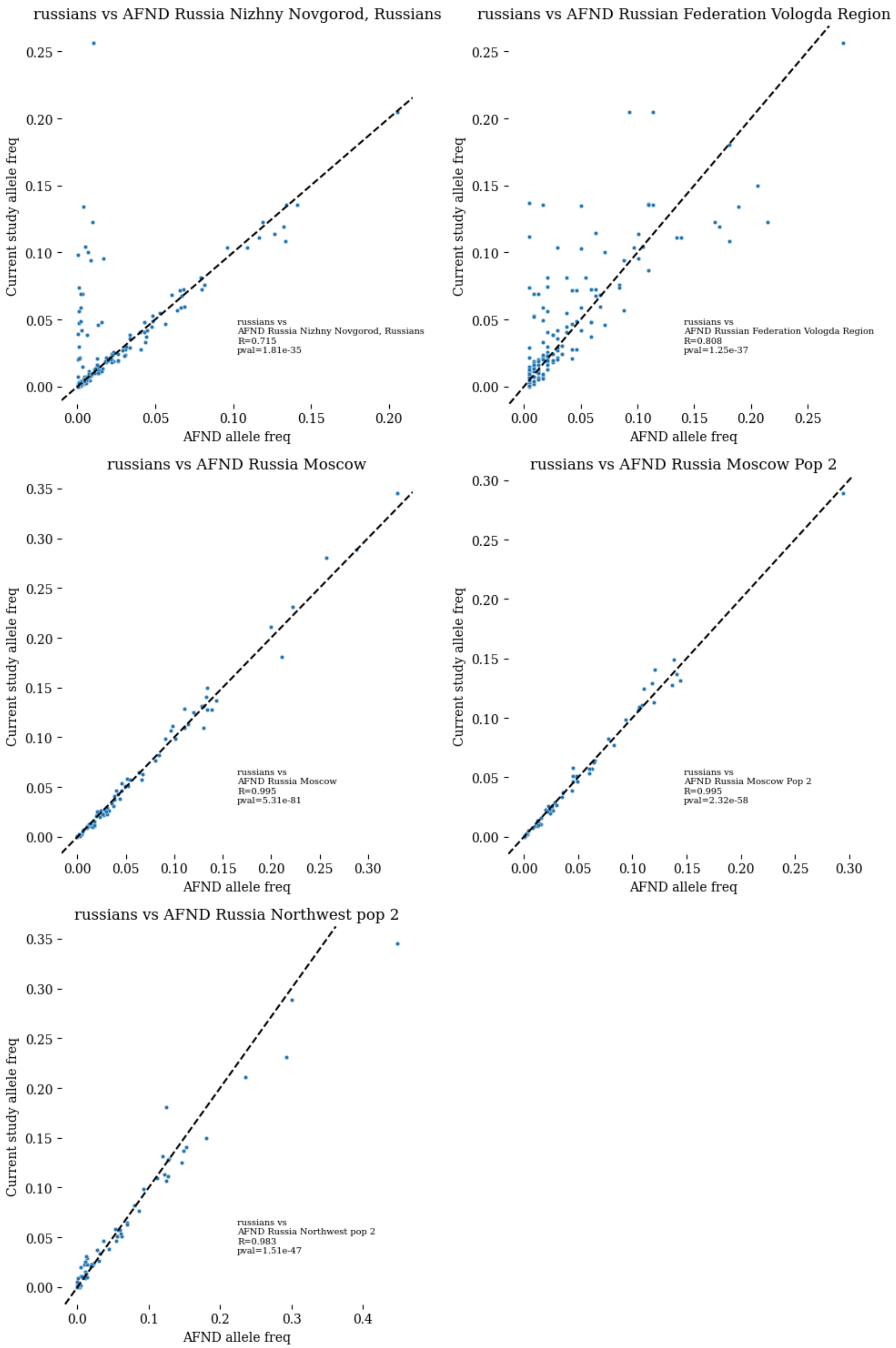


Fig S5. Allele frequency correlation of the current study cluster russians and the corresponding AFND datasets, with the black dashed line representing the line of perfect haplotype frequency agreement (y=x).

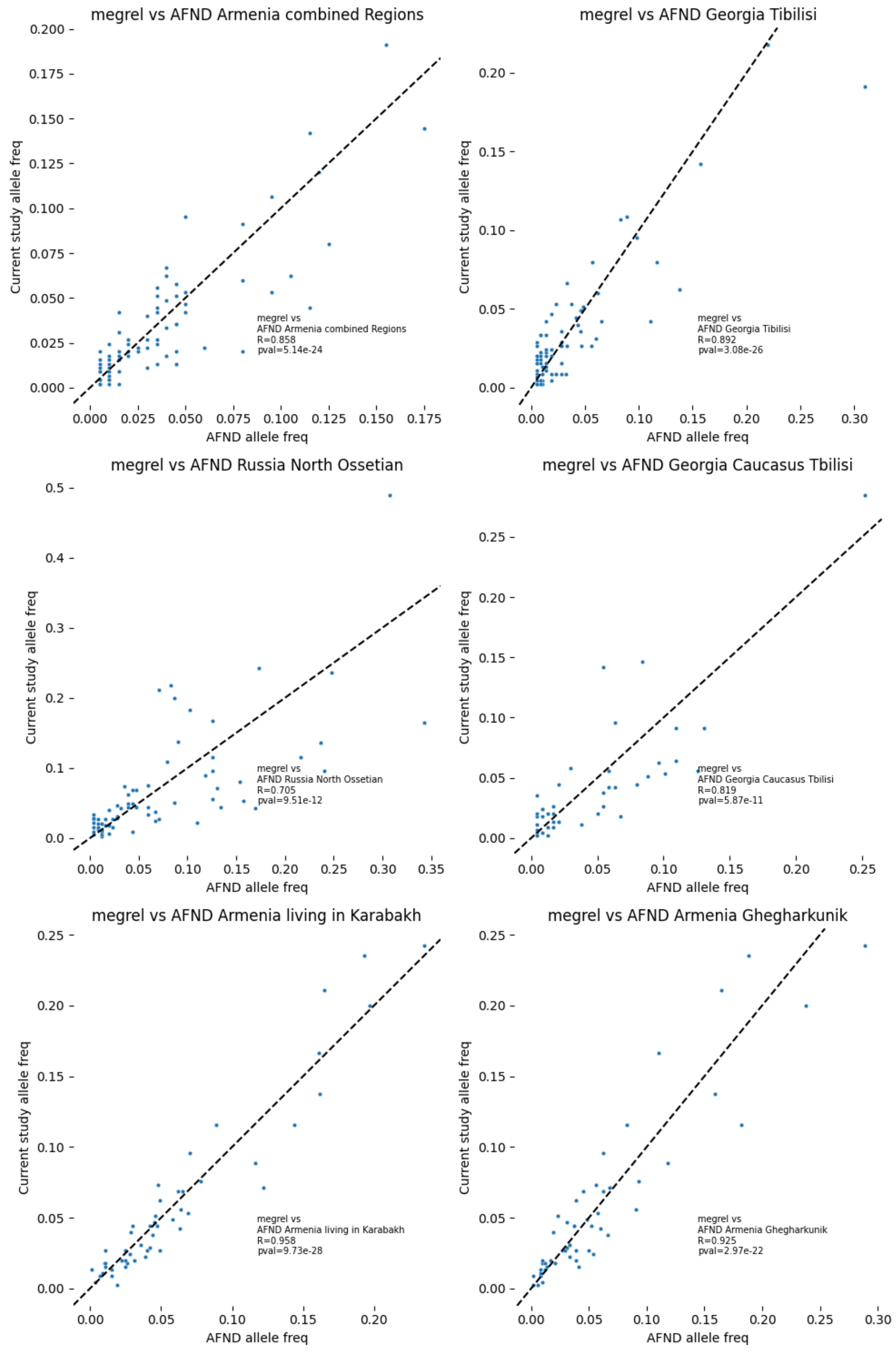


Fig S6. Allele frequency correlation of the current study cluster megrel and the corresponding AFND datasets, with the black dashed line representing the line of perfect haplotype frequency agreement (y=x).

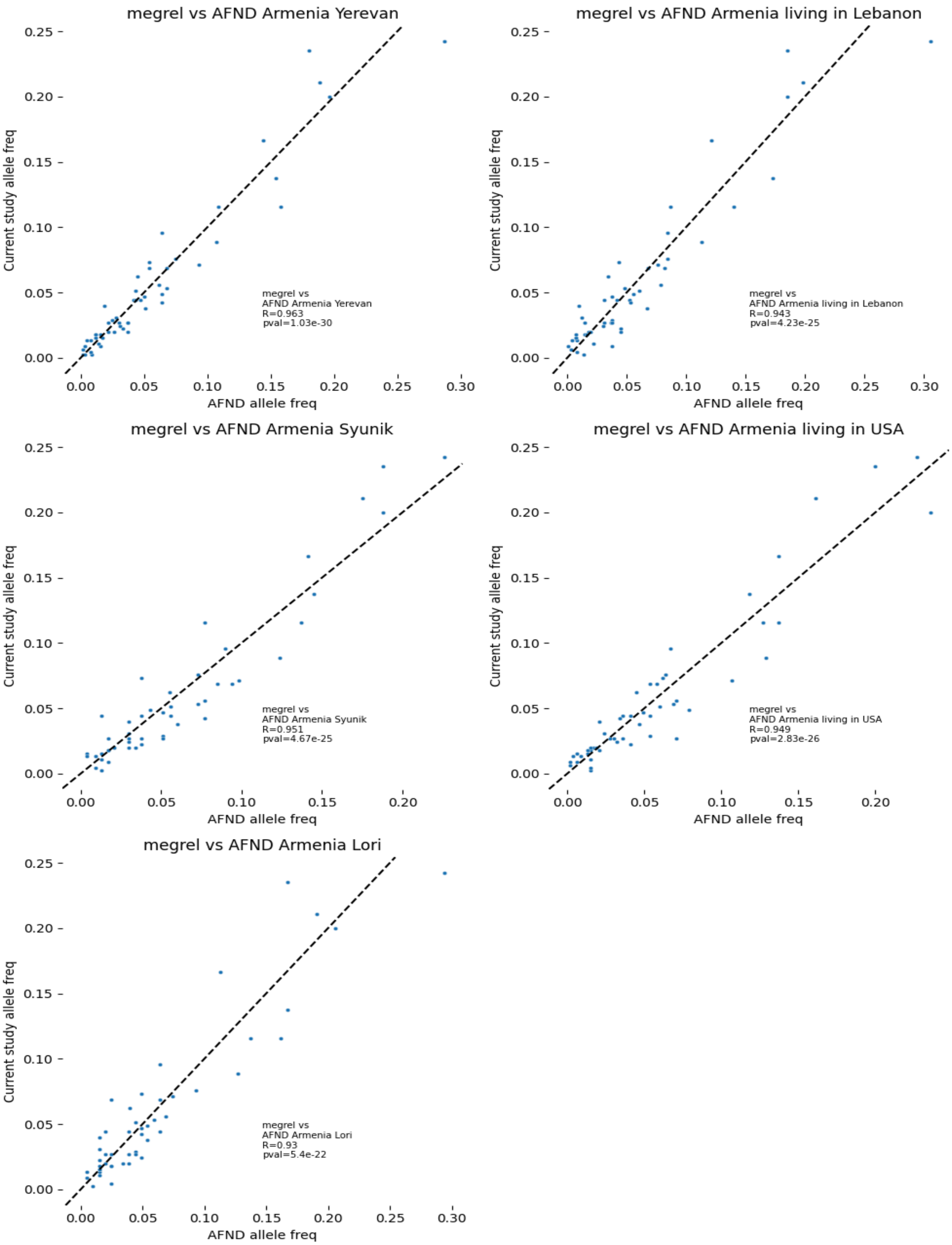
megrel vs AFND Armenia Yerevan
megrel vs
AFND Armenia Yerevan
R=0.963
pval=1.03e-30
megrel vs AFND Armenia living in Lebanon
megrel vs
AFND Armenia living in Lebanon
R=0.943
pval=4.23e-25
megrel vs AFND Armenia Syunik
megrel vs
AFND Armenia Syunik
R=0.951
pval=4.67e-25
megrel vs AFND Armenia living in USA
megrel vs
AFND Armenia living in USA
R=0.949
pval=2.83e-26
megrel vs AFND Armenia Lori
megrel vs
AFND Armenia Lori
R=0.93
pval=5.4e-22
Current study allele freq
AFND allele freq

Fig S7. Allele frequency correlation of the current study cluster megrel and the corresponding AFND datasets, with the black dashed line representing the line of perfect haplotype frequency agreement (y=x).

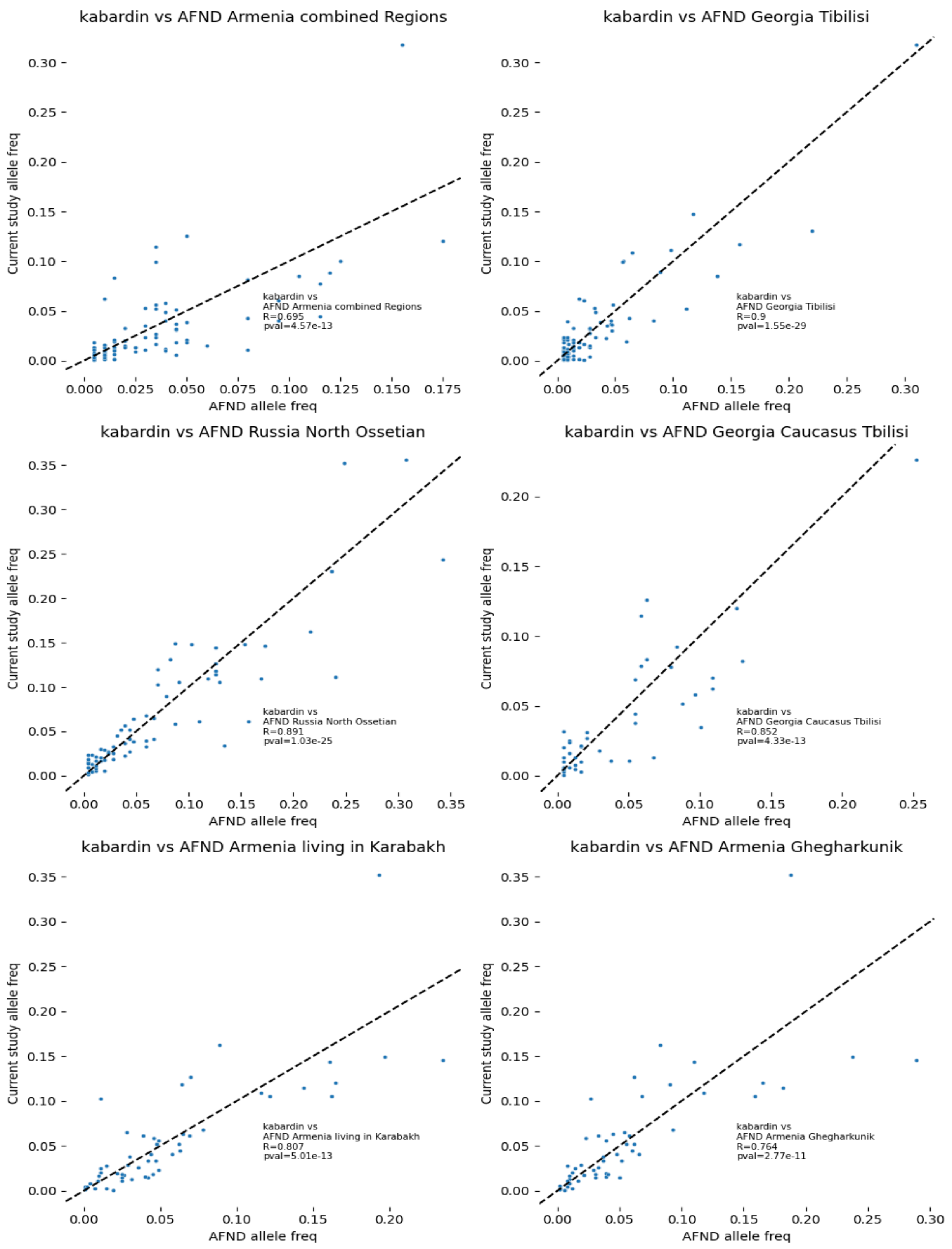
kabardin vs AFND Armenia combined Regions
kabardin vs
AFND Armenia combined Regions
R=0.695
pval=4.57e-13
kabardin vs AFND Georgia Tibilisi
kabardin vs
AFND Georgia Tibilisi
R=0.9
pval=1.55e-29
kabardin vs AFND Russia North Ossetian
kabardin vs
AFND Russia North Ossetian
R=0.891
pval=1.03e-25
kabardin vs AFND Georgia Caucasus Tbilisi
kabardin vs
AFND Georgia Caucasus Tbilisi
R=0.852
pval=4.33e-13
kabardin vs AFND Armenia living in Karabakh
kabardin vs
AFND Armenia living in Karabakh
R=0.807
pval=5.01e-13
kabardin vs AFND Armenia Ghegharkunik
kabardin vs
AFND Armenia Ghegharkunik
R=0.764
pval=2.77e-11
Current study allele freq
AFND allele freq

Fig S8. Allele frequency correlation of the current study cluster kabardin and the corresponding AFND datasets, with the black dashed line representing the line of perfect haplotype frequency agreement (y=x).

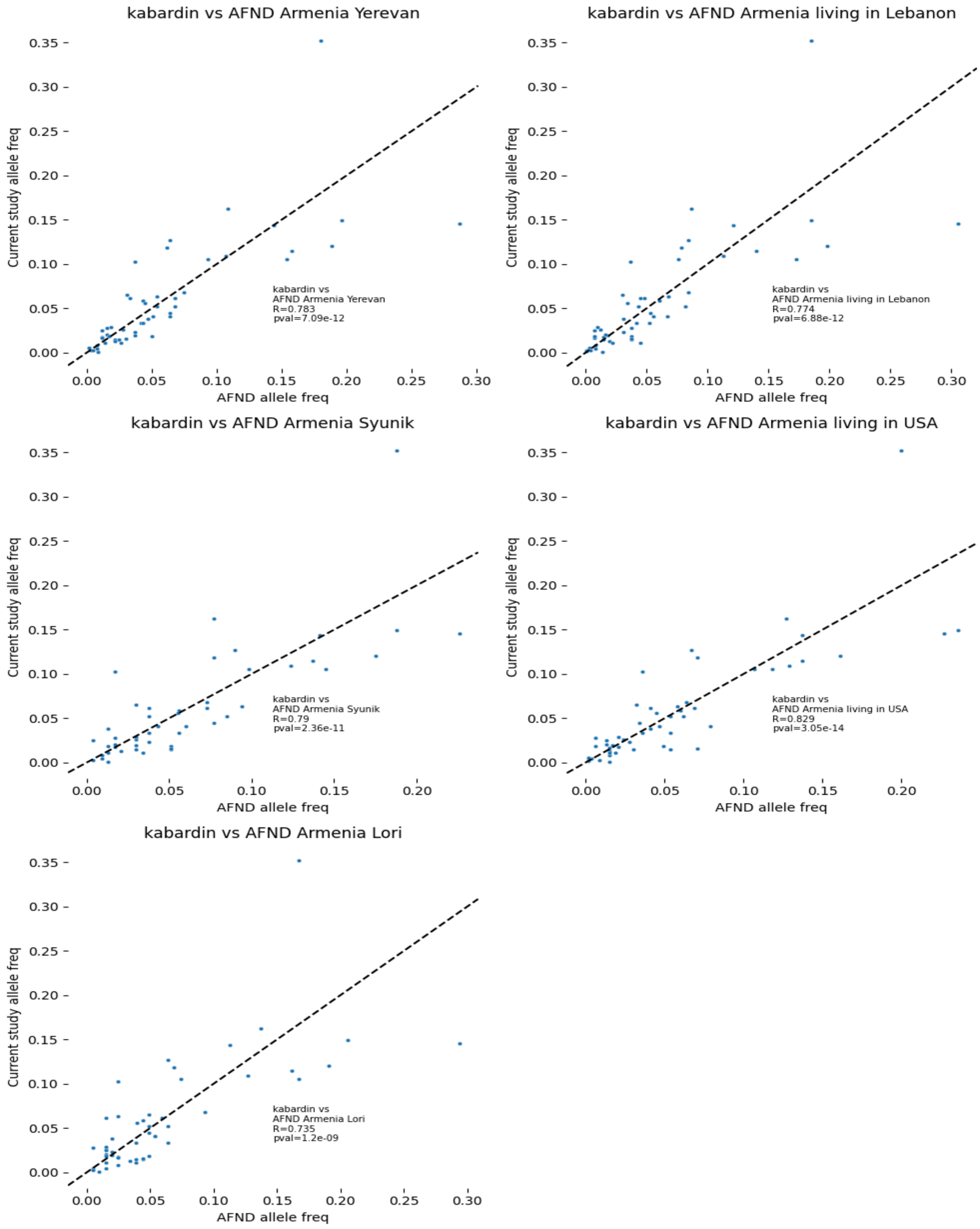

Fig S9. Allele frequency correlation of the current study cluster kabardin and the corresponding AFND datasets, with the black dashed line representing the line of perfect haplotype frequency agreement (y=x).

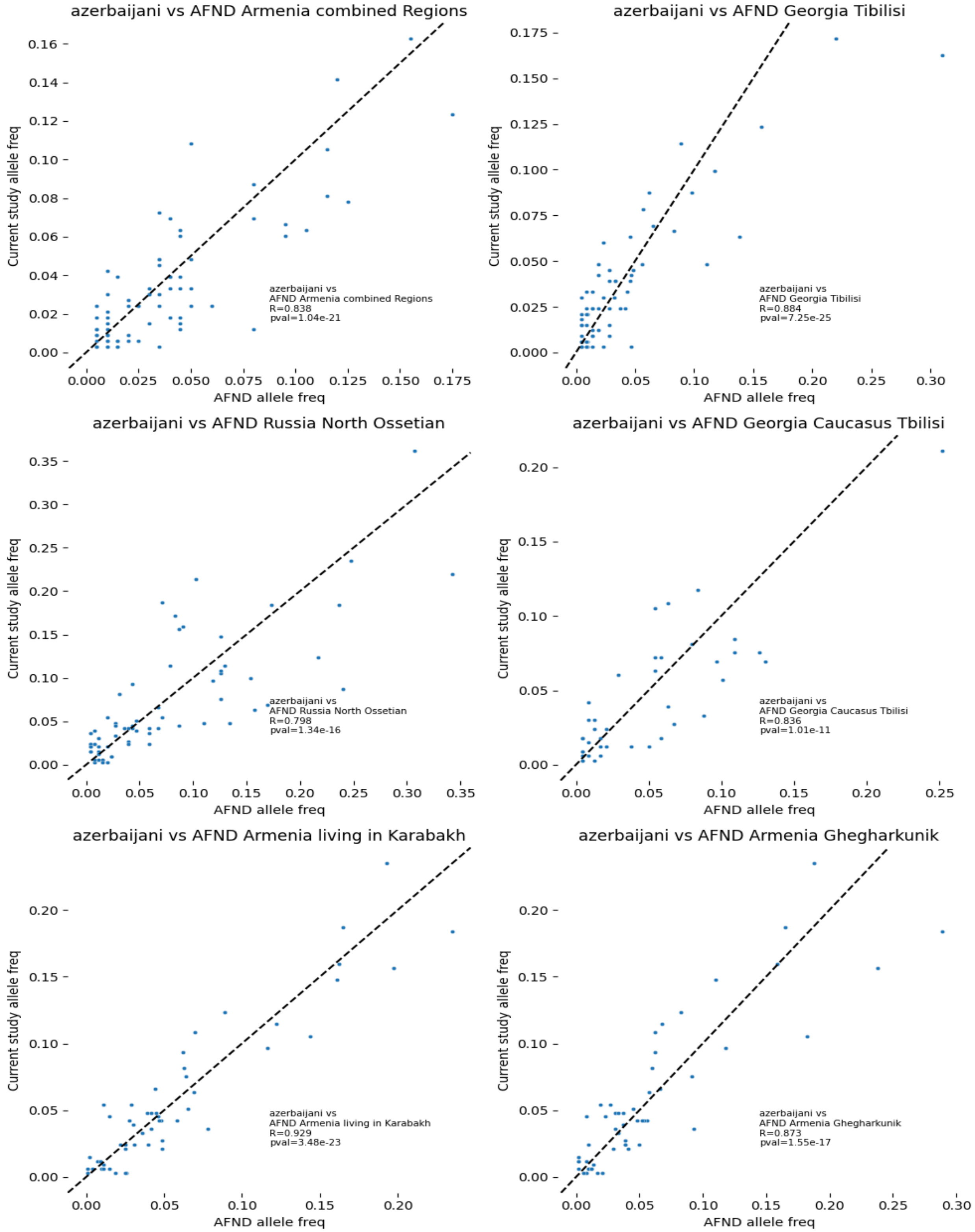

Fig S10. Allele frequency correlation of the current study cluster azerbaijani and the corresponding AFND datasets, with the black dashed line representing the line of perfect haplotype frequency agreement (y=x).

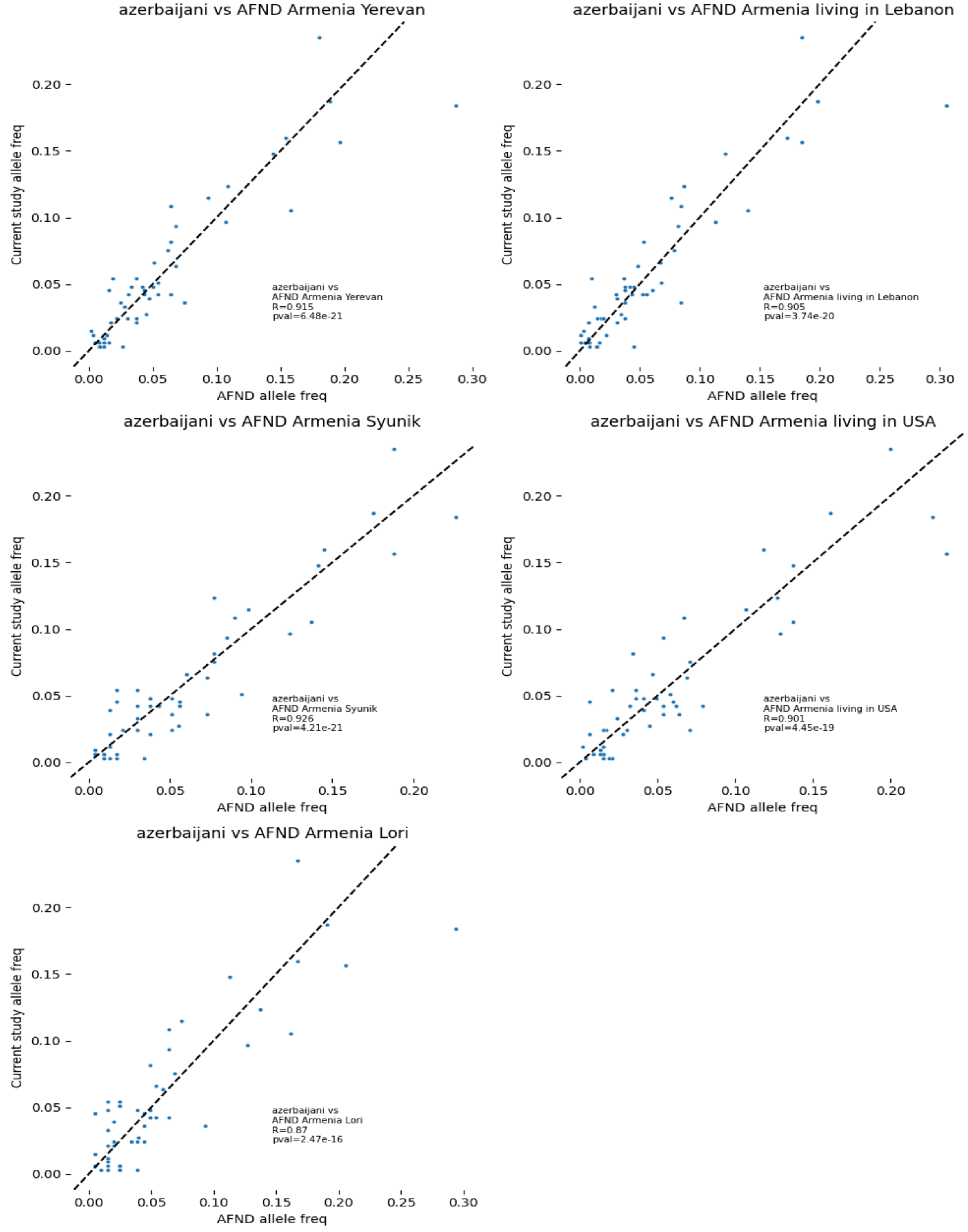

Fig S11. Allele frequency correlation of the current study cluster azerbaijani and the corresponding AFND datasets, with the black dashed line representing the line of perfect haplotype frequency agreement (y=x).

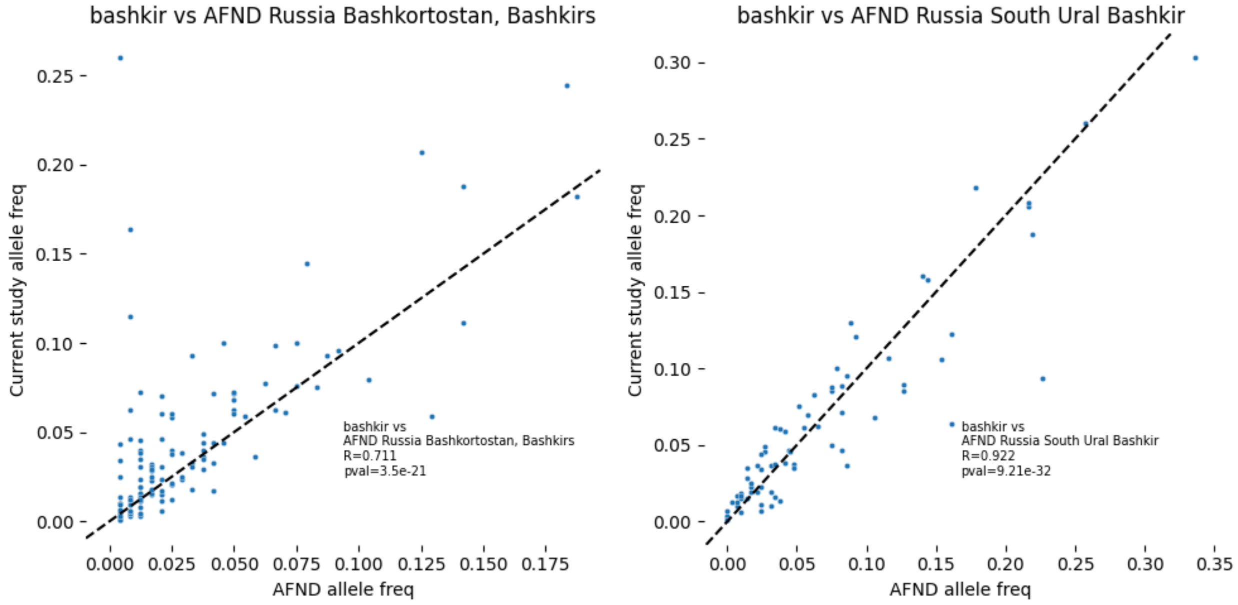


Fig S12. Allele frequency correlation of the current study cluster bashkir and the corresponding AFND datasets, with the black dashed line representing the line of perfect haplotype frequency agreement (y=x).

Table S3. Summarized Pearson correlation coefficients for pairwise allele frequency comparisons between the study ethnic clusters and populations from the AFND. Values represent the mean, median, and standard deviation (SD) of correlation coefficients calculated across all valid population pairs within each cluster.

| | **Mean** | **Median** | **SD** |
|---|---|---|---|
| **azerbaijani** | 0.879567 | 0.884431 | 0.041635 |
| **bashkir** | 0.816571 | 0.816571 | 0.148789 |
| **buryat** | 0.832332 | 0.78839 | 0.087048 |
| **kabardin** | 0.801889 | 0.790426 | 0.062916 |
| **megrel** | 0.899475 | 0.929512 | 0.078776 |
| **russians** | 0.899241 | 0.983245 | 0.129977 |
| **tatars** | 0.80247 | 0.956707 | 0.290810 |
| **tuva** | 0.867607 | 0.864533 | 0.042541 |

Table S4. Population cluster pairs with significantly different distributions of T1D PRS values

| combination | pval | adjusted_pval |
|---|---|---|
| (yakut, russians) | 2.46E-21 | 2.24E-19 |
| (yakut, kabardin) | 2.75E-18 | 2.48E-16 |
| (yakut, tatars) | 6.65E-14 | 5.91E-12 |
| (kazakh, kabardin) | 1.55E-12 | 1.37E-10 |
| (kazakh, russians) | 3.62E-12 | 3.15E-10 |
| (buryat, russians) | 5.37E-11 | 4.62E-09 |
| (buryat, kabardin) | 6.58E-11 | 5.59E-09 |
| (yakut, bashkir) | 2.46E-10 | 2.06E-08 |
| (tuva, kabardin) | 3.47E-09 | 2.88E-07 |
| (tuva, russians) | 5.80E-09 | 4.75E-07 |
| (yakut, megrel) | 3.24E-08 | 2.63E-06 |
| (buryat, tatars) | 2.45E-07 | 1.96E-05 |
| (tatars, kabardin) | 3.05E-07 | 2.41E-05 |
| (kazakh, tatars) | 7.79E-07 | 6.08E-05 |
| (uzbek, kabardin) | 1.11E-06 | 8.56E-05 |
| (tatars, russians) | 3.58E-06 | 2.72E-04 |
| (tuva, tatars) | 4.16E-06 | 3.12E-04 |
| (buryat, bashkir) | 4.27E-06 | 3.16E-04 |
| (uzbek, russians) | 1.38E-05 | 1.01E-03 |
| (yakut, azerbaijani) | 2.03E-05 | 1.46E-03 |
| (tuva, bashkir) | 2.31E-05 | 1.64E-03 |
| (buryat, megrel) | 3.62E-05 | 2.53E-03 |
| (kazakh, bashkir) | 4.15E-05 | 2.87E-03 |
| (kalmyk, kabardin) | 9.03E-05 | 6.14E-03 |
| (tuva, megrel) | 9.68E-05 | 6.48E-03 |
| (yakut, mansi) | 1.79E-04 | 1.18E-02 |
| (kalmyk, russians) | 1.98E-04 | 1.29E-02 |
| (bashkir, kabardin) | 2.97E-04 | 1.90E-02 |
| (kazakh, megrel) | 3.77E-04 | 2.37E-02 |
